# Social media posts as a source of ecological information over time: using Twitter (X) as a proof of principle

**DOI:** 10.1101/2025.11.21.689797

**Authors:** Rafał Miłodrowski, Cassandra G Extavour, Guillem Ylla

## Abstract

Acquiring data about ecology over long periods of time and large geographical areas is often difficult, expensive, and takes a long time. Here, we explore whether, through millions of daily posts on social networks, we could be constantly and unconsciously accumulating data regarding important biological patterns over the years.

Data analysis from generalist social networks has been successfully applied in different research fields, most notably in political science and epidemiology. Here, we evaluate their potential and drawbacks for studying biological patterns of other organisms over time. To that end, we used millions of posts on Twitter (currently X), over 11 years, on four different insect groups: cockroaches, crickets, Monarch butterflies, and mosquitoes.

Our results show that through millions of daily tweets, we could identify temporal periodic patterns that reflect their unique known phenology of the four insect taxa studied. Given the 11-year span of the dataset, we could also track changes in their patterns over the years that might be related to environmental factors. Using a sentiment analysis, we could also characterize the emotions of people towards these animals, which is important for the design of awareness campaigns. We also discuss the limitations of these data, such as the potential that social media data might have for spatial tracking of migrations.

In summary, we show compelling evidence for the use of social network data for biologists and provide a theoretical and practical framework for exploring these sources of data.

## 1. Introduction

Acquiring data about ecology over long periods of time and large geographical areas is difficult and often expensive. Compiling information on geographical species distribution, animal behavior, or the impact of climate change on different animals requires costly expeditions, expensive equipment, and time. Additionally, data collected in these expeditions usually represent snapshots of a given time point, limiting our ability to observe changes over time.

What if we were constantly and unconsciously accumulating data regarding important aspects of ecology? That could be immensely useful. In this article, we test the hypothesis that through the millions of daily posts on generalist social media platforms, we are indirectly accumulating data on ecological patterns over time that can be abundant and reliable enough to be useful for research and conservation. For that purpose, we use Twitter posts (currently X) referring to four insects covering the years 2011 to 2022.

Social networks developed from a tool used by a small group of people to what it is today: a channel of communication and interaction used by millions of people worldwide^1^. In about two decades, social media platforms grew from simple means of communication to diverse and complex systems, allowing anonymous individuals, also institutions, professionals, politicians, and scientists to communicate on a global scale^2^.

Among the current most popular social networks is Twitter (currently named X), a free social networking site where users broadcast short posts known as tweets, and which has become particularly relevant in many aspects of today’s reality: from instant news to public engagement with science and politics. Political personalities have used Twitter as one of the most important media for direct access to the public, public opinion, and mobilization of electorates^3,4^. The political uses of Twitter extend well beyond election campaigns, with the potential to organize protests and mobilize masses, as evidenced by the Arab Spring, Black Lives Matter, and global climate strikes^5^. This has given rise to the phenomenon now referred to as "networked protest," in which social movements can plan and coordinate activities across the globe without the need for a central organization entity^6^. The speed and accessibility of Twitter enable these movements to spread messages further than ever before, often resulting in significant social and political impacts over a short period. However, Twitter’s role in political activism also has its negative aspects, such as the spread of fake news and polarizing debates^7^.

In addition to its political uses, Twitter has also proved to be a useful tool in scientific research in different disciplines, such as in epidemiology^8^, climate change^9^, and public health^10^. Remarkably, during the COVID-19 pandemic, Twitter was used as a tool for real-time monitoring of the pandemic, public health information dissemination, tracking misinformation flow, as well as understanding public perceptions regarding vaccination programs and government policies^11^. Twitter has also been used by epidemiological modeling and communications strategies for other diseases, such as in detecting potential depression through sentiment analysis in both English and Arabic tweets^12^, or determining the perceptions of different contraceptive methods^13^.

Beyond public health, scientists have used Twitter to assess public interest in environmental and biodiversity issues. Pew Research Center (2019)^14^ explored public engagement with topics like climate change, conservation, and the loss of biodiversity, showing that while high-visibility topics such as global warming consistently generate substantial traffic, less popular subjects, such as the conservation of insects and arthropods, also receive significant attention. To study how the conservation status of species correlates with public awareness and research efforts, Wang et al. (2021)^15^ used multiple resources on the Internet, including 500 tweets per month. This research revealed that neither public nor research interests are correlated with the conservation status of species, highlighting a worrying neglect of certain endangered animal taxa.

Sharing experiences in writing is not exclusive to social networks. Humans have done it for centuries and in many forms, among which are the haiku (short Japanese poems). For example a study of 4,000 haiku poems, unveiled temporal trends, perceptions, and emotions of people about arthropod species^16^. Nowadays, in contrast with publishing 17th-century haiku, everyone from any part of the world can share their experiences using social networks, making them a potentially very powerful tool for understanding temporal and geographical trends and emotions about specific topics.

Although everyone can share their experiences on these networks, the platform’s algorithms might prioritize specific topics, which can skew public attention toward sensationalized or popular issues, while critical but less visible topics may be ignored^17^. This misalignment can also affect research priorities, funding, and policy decisions, which might have a long-lasting and profound effect on society. For instance, it has been shown that biases in media spread through the internet have strongly influenced schoolchildren’s perception of biodiversity, leading them to prioritize conservation efforts of exotic species over local species that are less represented in media^18^.

For ecology, social media has been proposed as a potential tool for citizen science projects^19^ and for monitoring species distributions^20,21^. Building on this idea, another study obtained a manually-curated dataset of 8,698 tweets from periods of ∼6 months across four years in the UK containing specific hashtags related to ants, spiders, and starlings, with which they showed it could reproduce the temporal ecological patterns obtained through three citizen-science projects^22^.

Here, we explore the challenges and opportunities that social networks offer for ecological research and for the understanding of people’s emotions about biodiversity. For that purpose, we obtained 1.6 million tweets from Twitter posted over 11 years on four insect groups, as a means of evaluating the potential power and caveats of social network posts as a reliable source of information for research in ecology and conservation biology.

## 2. Results

### 2.1. Data Description

To check whether social networks might contain biologically relevant information, we decided to use Twitter (currently X) as proof of principle, due to its high volume of users and posts (called Tweets) at the time of our analysis (with circa 450 million users and 500 million daily posts in 2023^23^), the fact that it had been previously used for academic purposes as described above, and for which we obtained an academic license for API access to perform the four queries of interest on the whole database. We queried posts of four different abundant and commonly recognized insects with independently well-characterized and distinct behavioral patterns: Monarch butterflies, mosquitoes, crickets, and cockroaches.

Using the Twitter API for academics, we downloaded a total of 154,009 tweets for cockroaches, 62,289 for crickets, 350,066 for Monarch butterflies, and 1,043,571 for mosquitoes, posted between 2011 and 2022 (**Table 1**). For each Tweet, in addition to the text of the tweet itself, we obtained additional data, as follows: unique identifier of the posting user, UTC datetime that the tweet was created, the unique identifier of the requested tweet, the geolocation of the tweet, the user name, and the user profile location.

**Table 1:** Description of the four datasets. For each insect query, the first row contains the phrase used to search the Twitter database, which excludes retweets and replies, and selects those classified as written in English. The second row shows the total number of Tweets retrieved with the query. The third row shows the number of tweets geo-data, while the fourth row shows the total number of tweets with geo-data plus those with the user’s location. The final row displays the number of tweets used in the analysis, which are those with either geo-data or geo-data from the user, after removing key words and selecting only those from the USA and Canada. Values in brackets indicate the percentage over the original number of tweets downloaded.

|  |  | **Cockroach** | **Cricket** | **Monarch butterfly** | **Mosquito** |
| --- | --- | --- | --- | --- | --- |
| **1** | **Search phrase** | '(found OR saw)<br>(cockroach OR cockroaches) -is:retweet -is:reply lang:en | '(crickets OR cricket) chirp -is:retweet -is:reply lang:en | 'monarch butterfly -is:retweet -is:reply lang:en | 'mosquito bite -is:retweet -is:reply lang:en |
| **2** | **Number of Tweets downloaded** | 154,009 | 62,289 | 350,066 | 1,043,571 |
| **3** | **Tweets with geo-data coordinates** | 4,937 (3.20%) | 1,671 (2.68%) | 9,615 (2.75%) | 31,972 (3.06%) |
| **4** | **Tweets with geo-data data including user's location** | 83,707 (54.35%) | 33,327 (53.50%) | 242,867 (69.37%) | 533,090 (51.08%) |
| **5** | **Tweets after filtering** | 36,413 (23.64%) | 17,964 (28.84%) | 145,528 (41.57%) | 286,108 (27.42%) |

A particular piece of information that could be relevant to scientists is the geolocation of the posts, namely the GPS coordinates at the time of posting, which could be an indicator of the location of a given observation. This geolocation of the tweet, is an option that the user must choose to activate while posting, and was only present in around 3% of tweets obtained (**Table 1**). A potential proxy for the location of the tweet could be the location that the user displays in their profile. This can be usually set by the user in terms of a place name (for example, the name of a University).

Using the OpenStreetMap^24^ with tidygeocoder^25^ R package, we obtained the coordinates of the centroid of the location name for 55% of the users. Combining both sources of location (geolocation of the tweet and self-reported location of the user), circa 58% of the tweets could be linked to coordinates (**Table 1**).

Plotting the raw number of tweets retrieved per query per year revealed a distinct annual repeating pattern for each query (**Figure 1**). The fact that we observed consistent, distinct annual patterns for each query, suggested we were observing not only noise of the data, but specific recurring patterns for each queried insect group that could reflect their biological patterns. Despite observing a clear temporal signal overall, we attempted to remove the main potential sources of noise by removing those from shops that contained the words “etsy”, ”ebay”, or “store”, and retaining only tweets for which either the tweet or the user was located in the USA and Canada, to geographically delimit the scope of our analysis. This retained 23.64% of tweets from cockroaches, 28.84% from crickets, 41.57% from Monarch butterflies, and 27.42% from mosquitoes (**Table 1**).

**Figure 1:**
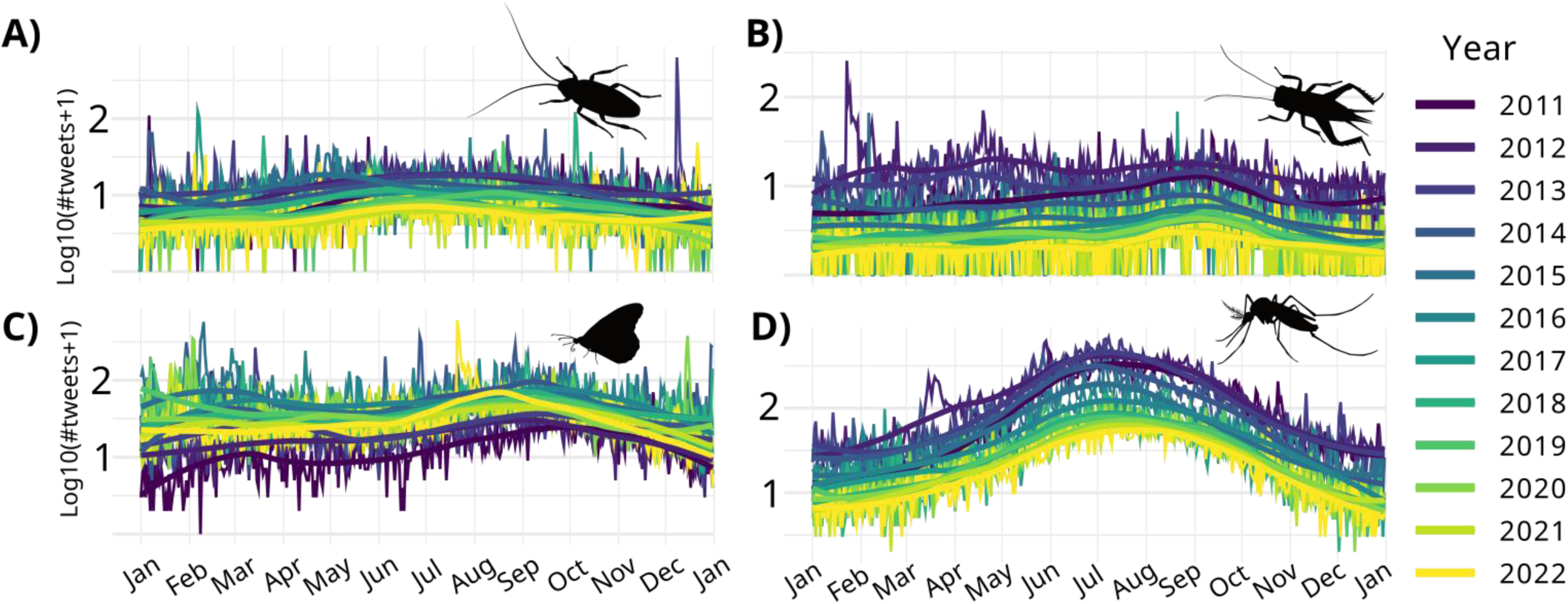
The number of tweets (in log10 scale) and the smoothen line (loess function) along the 2011-2022 (by color) for A) cockroaches, B) crickets, C) monarch butterfly and D) mosquitoes, reveal a distinct pattern for each insect group that repeats yearly.

### 2.2. Seasonal Patterns from Twitter data

With the filtered Tweets from the USA and Canada, we proceeded to ask whether this dataset could provide information about the seasonal patterns of these species over time. For that purpose, we first computed the number of tweets per day of the year for each query across the 11 years. This revealed clearly different temporal patterns for each of the four queries (**Figure 2**). The trends became even more evident after applying a Temporal Analysis Using the X-11 Model, which subtracts the underlying trends and noise to highlight recurrent temporal patterns (**Figure 3 and Supplementary Figure S1-4)**.

**Figure 2:**
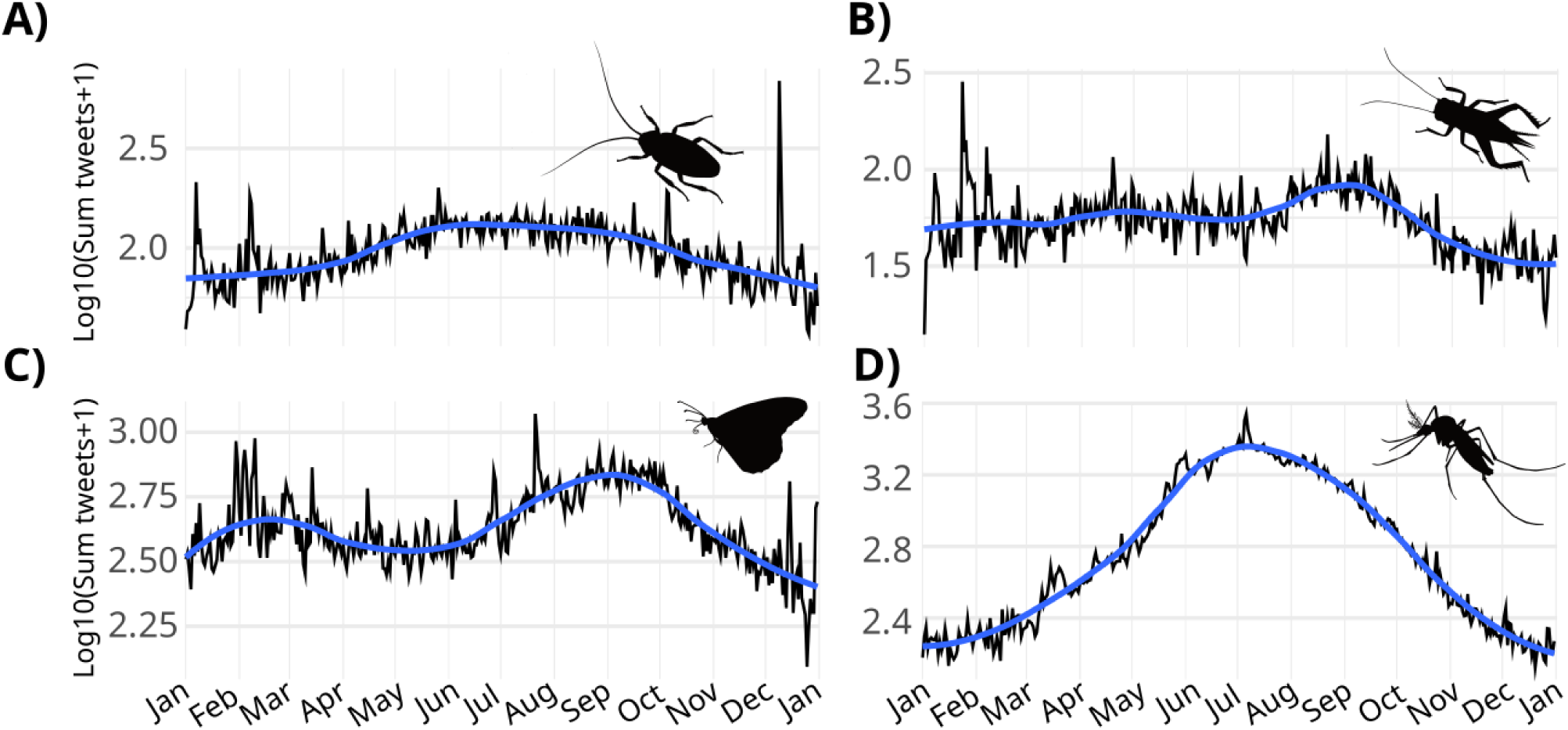
Sum of the number of tweets from the USA and Canada per day of the year (in log10 scale) and the smooth line (loess function) in blue for each query. D) The cockroach tweets are abundant throughout the year, but less frequent in the colder months (December to March). B) Cricket tweets peak in the summer months (August and September), during which they are more active and mating^27^. C) The monarch butterfly tweets display two peaks, one in early spring (February-April), and a second one in summer and fall (August-November), which may correspond with the migration northwards and then southwards^28^. D) Mosquito tweets are common throughout he summer (April to October), peaking in the warmest months (around July).

**Figure 3:**
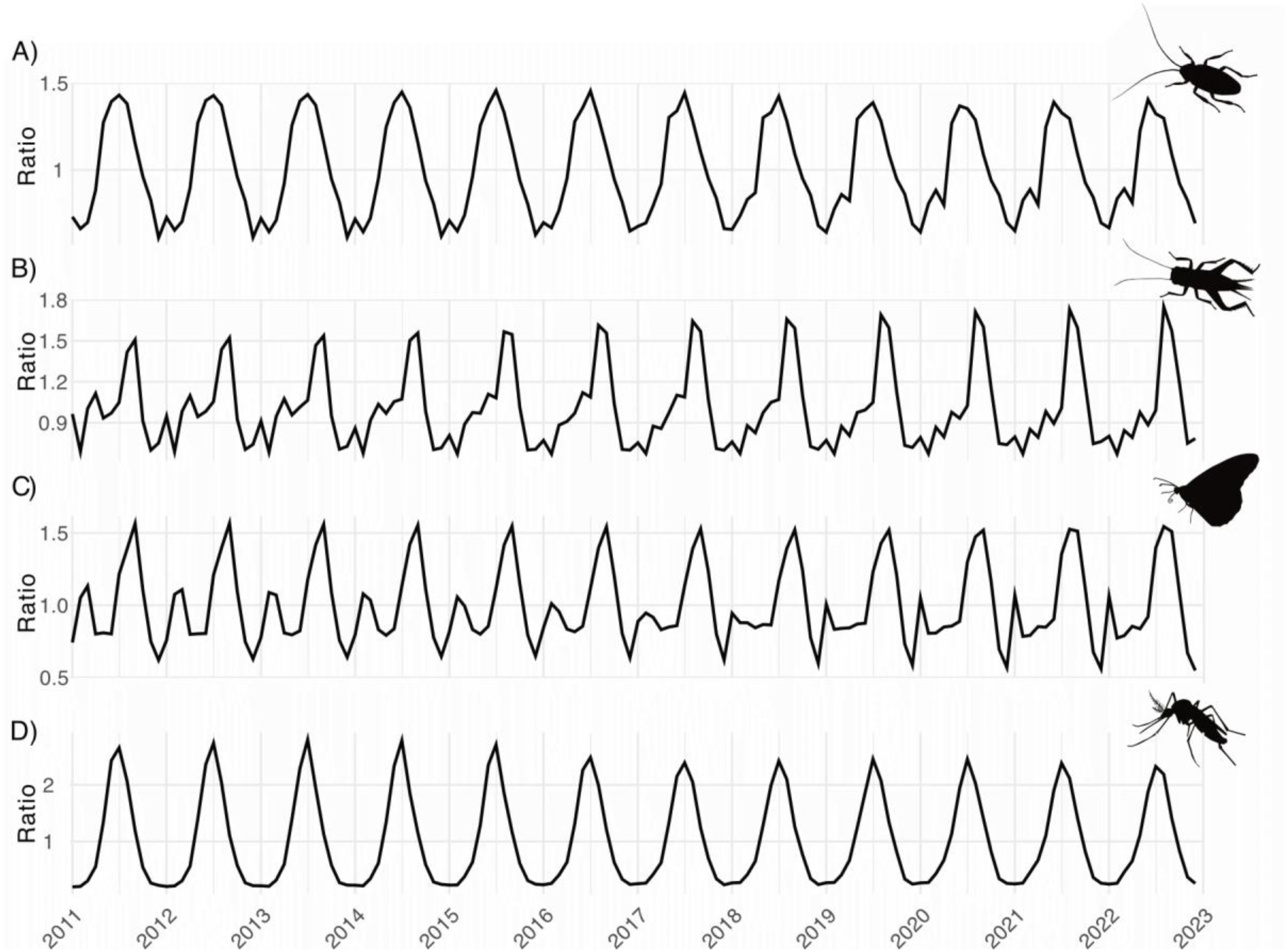
Seasonal decomposition done with X11 model highlights signal over 2011 and 2022 years in the number of tweets (as ratio to the annual mean centered at one) for A) cockroaches, B) crickets, C) monarch butterfly, and D) mosquitoes, highlighting the annually consistent patterns which are unique to each insect group.

Based on these analyses, we hypothesize that these seasonal profiles are not solely a reflection of users’ tweeting habits, but that they also reflect the observations that Twitter users make of these species, thus unveiling their biological patterns. These patterns suggest the insects’ activity over the year in the USA and Canada, matching the known activity patterns of these species: The cockroaches, being an indoor pest, are present year-round in homes but less abundant in the coldest months^26^. Crickets chirp in the late summer nights when it is warm outside and are ready to mate^27^. The characteristic Monarch butterflies with their migration northwards in early spring and southwards in the fall^28^. Mosquitoes, as it is well known, bite during the summer months^29^.

By contrast to the seasonal patterns, we did not identify any query-specific pattern based on the hour of the day or based on the day of the week. We therefore speculate that these patterns (lower number of tweets during night, and weekend) indeed reflect users’ activity patterns rather than insect activity patterns (**Supplementary Figure S5**).

### 2.3. Seasonal patterns in a specialized social network

We wondered whether and how the observations on temporal patterns from a generalist social network (like Twitter) compared with those of a specialized social network such as iNaturalist. For that purpose, we retrieved from the Global Biodiversity Information Facility (GBIF) the research-grade observations (those with media support, geolocation, and community consensus on the species identification) provided by iNaturalist for crickets (*Gryllidae*), cockroaches (*Blattodea*), the Monarch butterfly (*Danaus plexippus*), and mosquitoes (*Culicidae*), covering the same years 2011-2022 for the North America (in this case, the dataset also includes Mexico).

This resulted in 4,666 records for cockroaches, 17,988 for crickets, 142,292 for the Monarch butterfly, and 12,221 for mosquitoes. Compared to the Twitter dataset, this represented an order of magnitude fewer entries than tweets for cockroaches, almost the same number of entries for cricket, slightly fewer entries for the Monarch butterfly, and only 4% of the number of mosquito tweets.

As with Twitter datasets, we aggregated occurrences to monthly counts per query and used X-11 decomposition to extract the seasonal components (**Supplementary Figures S6-S9**). Despite obvious differences in both datasets (for example, in iNaturalist, there were observations of many different species of cockroaches and crickets, which are not necessarily the ones people would commonly see in their daily lives), the seasonal patterns were mostly in agreement with those obtained in the Twitter dataset for the last few years. However, we note that there were very few observations prior to 2020. One of the most remarkable differences between the two datasets, was that the spring peak for Monarch butterflies appeared lower in the iNaturalis dataset.

This comparison shows that a generalist social network, like Twitter, despite being very noisy, contains a signal that matches that of the expert research-grade observations posted in iNaturalist. Thus, while the number of observations in specialized databases might be more limited and skewed towards specific countries, generalist social networks might contain general trends that are equally valuable and might cover a larger and more diverse area, given that there are generalist social network users (not necessarily only on Twitter) around the globe.

### 2.4. Change in seasonal patterns

Given that our dataset contains information from 11 years, we asked whether it could be used to detect changes in the temporal patterns of the insects queried over those years, which could yield valuable information to study changes in biological patterns over time. Our analysis suggests that this is indeed possible (**Figure 4**).

**Figure 4:**
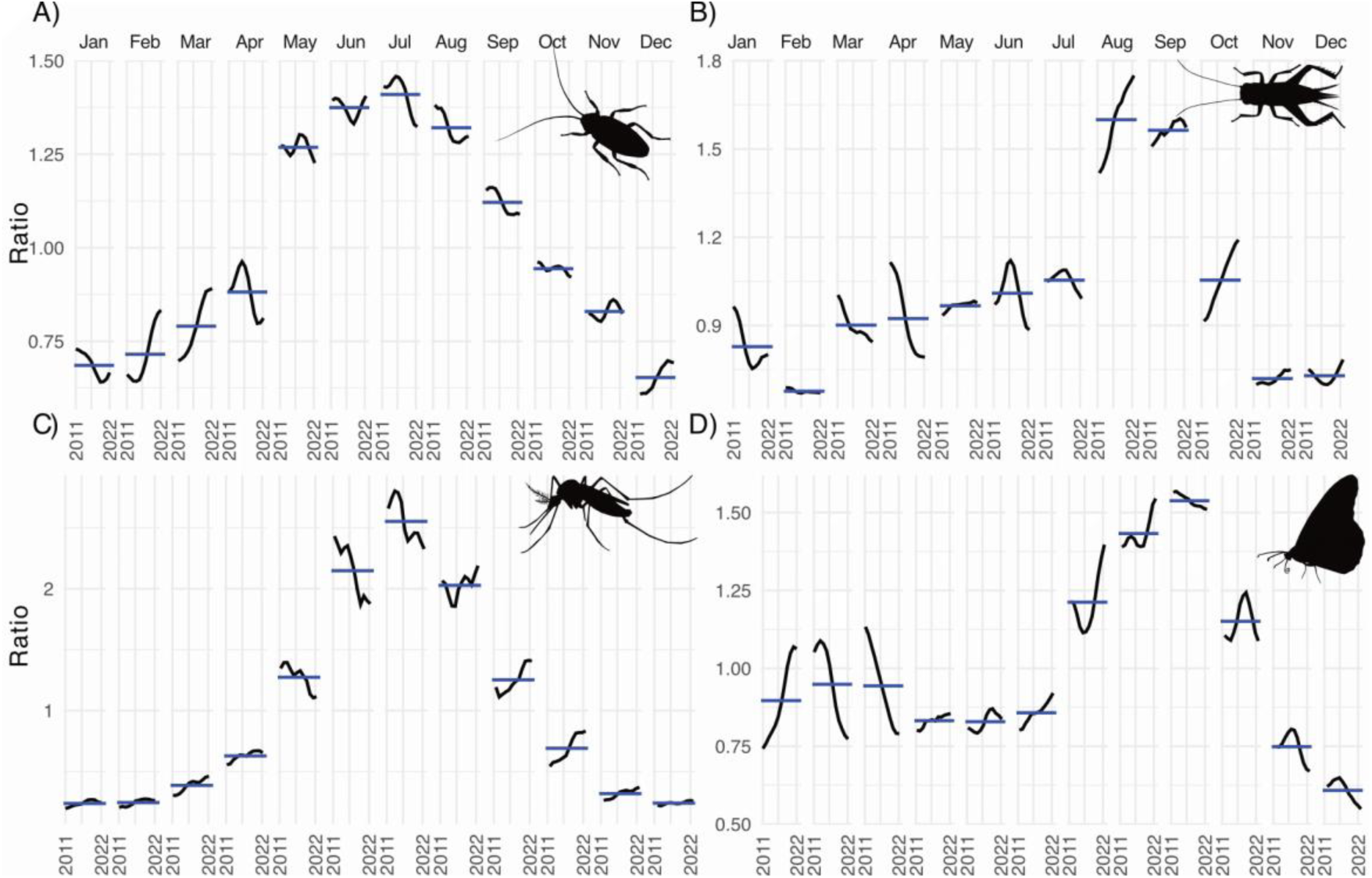
Seasonal changes in the ratios along the analyzed years on the monthly frequencies of tweets for A) cockroaches, B) crickets, C) monarch butterfly, and D) mosquitoes.

Cockroach tweets showed a tendency the years to increase their frequency earlier in the year (February-March), and they peaked a month earlier (June instead of July since 2020) (**Figure 4 & Supplementary Figure S10A**). Cricket tweets showed a tendency to advance the highest peak, thus shifting from September to August since 2015 (**Supplementary Figure S10B**) and extending the season towards the fall, with a clear increase in their frequency in the fall (**Figure 4**). The two peaks of tweets of Monarch butterfly showed a tendency to happen earlier over the years (**Figure 4 & Supplementary Figure S10C**). The mosquito peak appeared to occur in July over the years (**Supplementary Figure S10D**), while showing slightly lower frequency in June, and extending their longer in the season with increasing frequency over August, September, and October (**Figure 4**).

These findings suggest that social media posts (like tweets) could be used to identify changes in animal patterns over the years, which might in principle be linked to environmental and climatic data and changes. However, it should be noted that in this study, we are aggregating tweets from across the USA, a very large geographic area with different regional climates.

### 2.5. Emotional Distribution

Beyond research, dissemination is also an important aspect of conversation science. It is useful to know the feelings of people towards a given animal when planning conservation campaigns, educational activities, and policy-making. Hence, here we tested whether the social network posts could help us gain insights into the potential feelings of people towards specific animals using sentiment analysis and the presence of the most commonly used words.

Using the Emotion English DistilRoBERTa-base^30^ model from Hugging Face^31^ to perform emotion sentiment analysis on the text of the tweets, we assigned each tweet to the emotion with the highest score of the seven emotions included in the model (**Figure 5**). Here, we again observed very different patterns for each of the queried insect species.

**Figure 5:**
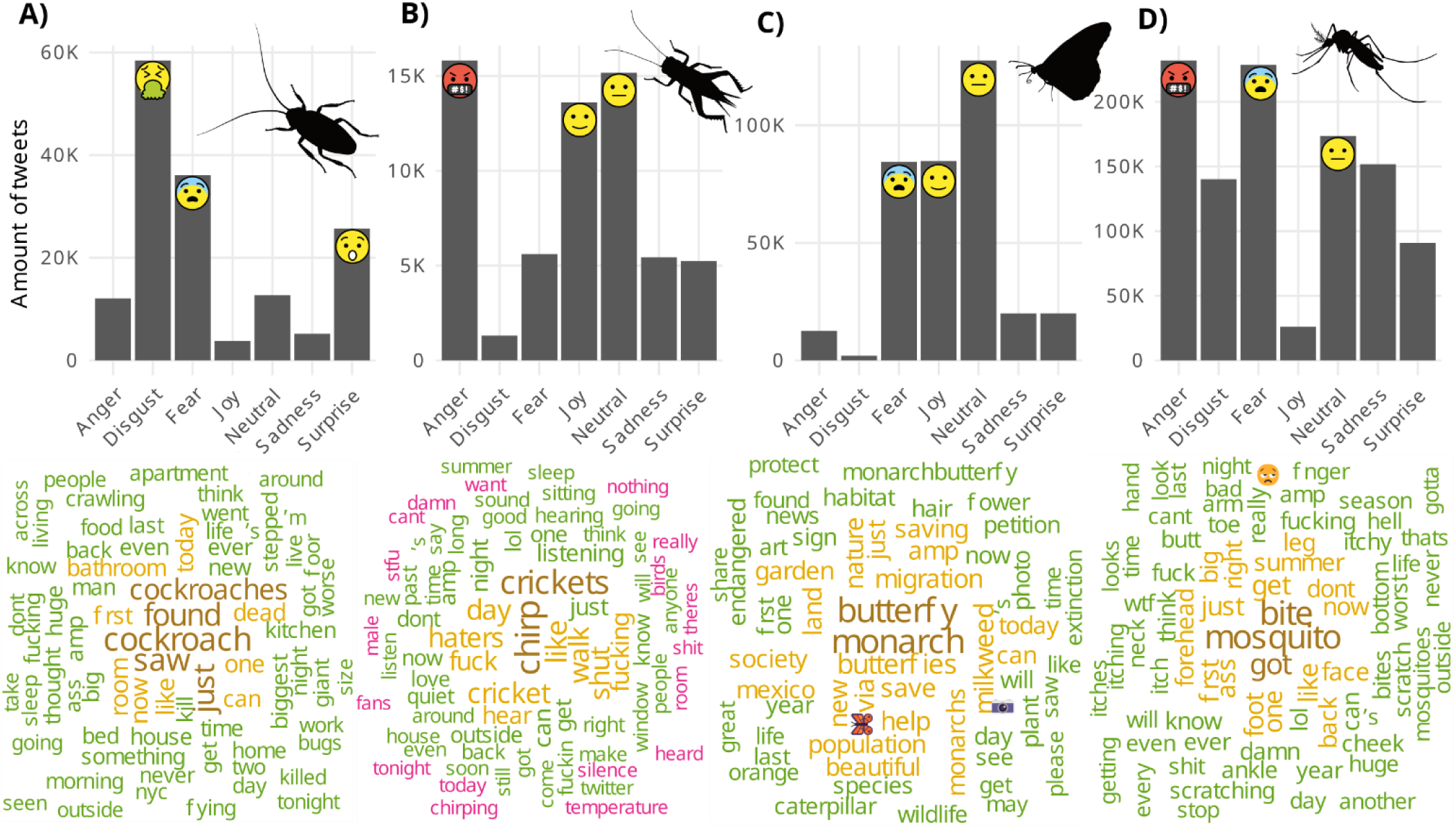
Number of tweets assigned to each sentiment (top) and word cloud (bottom) of the top 75 most frequently used words in the tweets about A) cockroaches, B) crickets, C) monarch butterfly, and D) mosquitoes. Top sentiments are highlighted with the corresponding emojis (from https://openmoji.org/).

The cockroach tweets, primarily reflected disgust, followed by fear, and surprise. The most frequent words often contained swear words, consistent with the described emotions and also information regarding where or when they had been most commonly spotted, including “bathroom” or “night”/”tonight”, which is consistent with the nocturnal habits on anthropic cockroaches and preference for humid indoor places^32^ **(Figure 5A**). Cricket tweets reflected primarily anger, joy, and neutral sentiments. The frequently used words suggested that anger comes from sleep deprivation due to their noisy chirping (**Figure 5B**). The monarch butterfly tweets were linked to feelings of fear, joy, and neutrality. We speculate that the fear may be due to their endangered status, supported by the observation of a high proportion of terms such as “help”, “save”, and “extinction” (**Figure 5C**). Mosquito tweets, were associated more often with anger and fear of their bites. The most frequently used terms not only expressed this anger, but also provided insights on the most commonly bitten body parts including “forehead”, “face“, “back”, “foot”, “leg”, and “ankle” (**Figure 5D**).

Overall, these analyses suggest that the text on the social media posts might also contain valuable information regarding the sentiments towards given species, as well as additional information, such as places where they are most often observed in the case of cockroaches, or where on the body they often bite people in the case of mosquitoes.

### 2.6. Geographical Distribution of Tweets

We further wondered whether the geolocation information could help track animal species, for example, to track the Monarch butterfly migration patterns over the years.

Using the geolocation of the tweet or that of the user profile, we analyzed geographical differences between the four queried insects (**Figure 6**). We observed, no clear differences between the four queries, all of which showed strong signals in regions with high population densities. Because most of the user locations were from a city or state, calculating the centroid created clusters of tweets in a specific city or point in the state. We also analyzed the tweets on the Monarch butterfly with geolocation data (not including those with location based on the user) by month of the year, to see whether this reflected the known migratory patterns and their change over time (**Supplementary Figure S10-S13**). However, we observed no clear trends. We speculate that this is due to the low number of tweets with geolocation data (2.75%), and the impact of highly populated areas. Even when attempting to normalize the signal by census data, we detected no clear trends distinct for each query.

**Figure 6:**
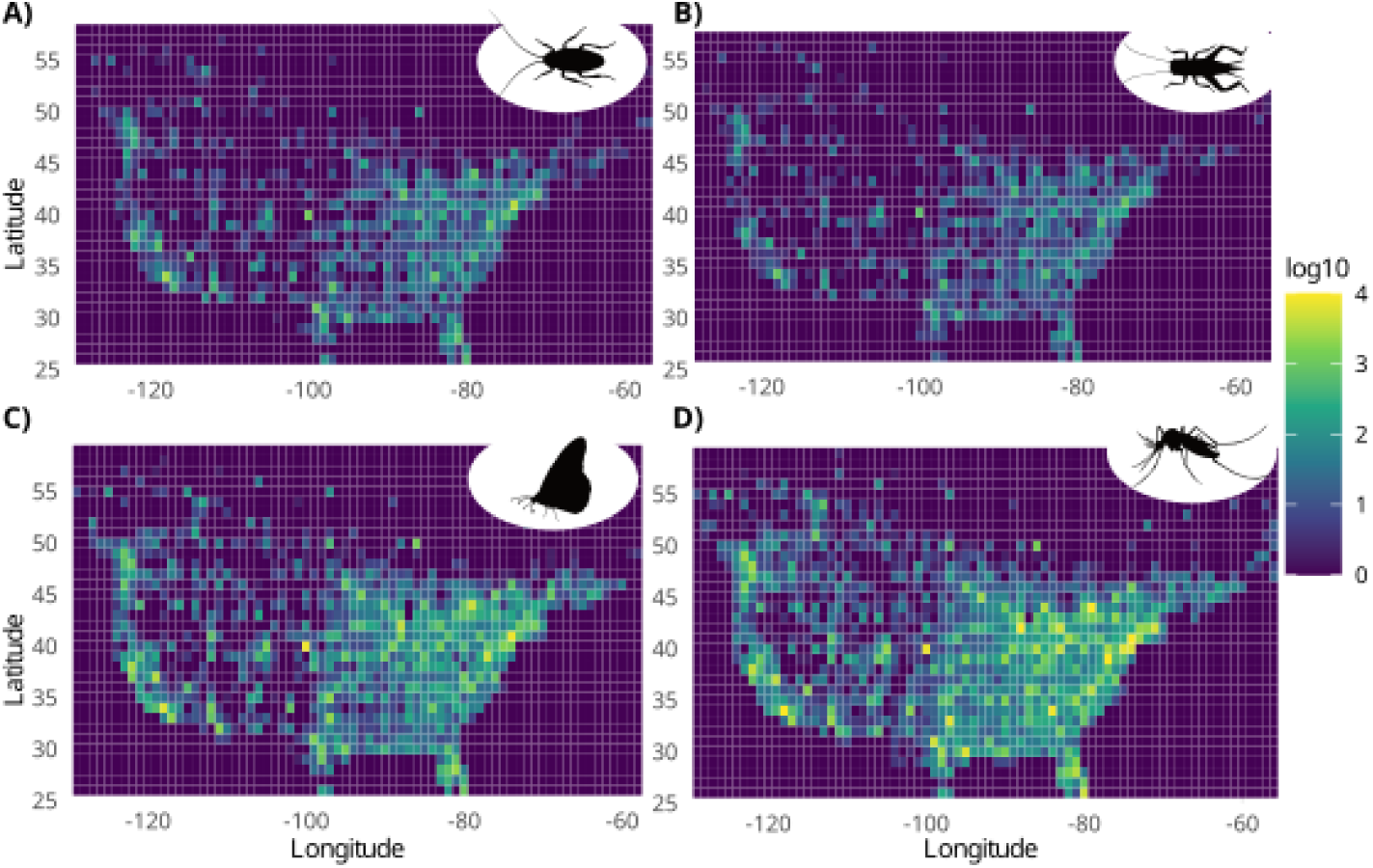
The location of the tweets in the USA and Canada, represented as a heatmap, shows the strong impact of highly populated areas. Color scale in represents the total number of tweets (log10 scale) for A) cockroaches, B) crickets, C) monarch butterfly, and D) mosquitoes in every square of one degree to one degree.

## 3. Discussion

### 3.1. Reliability of temporal patterns

Despite the large geographical area with heterogenous climate included in our analysis (USA and Canada), the temporal patterns observed in our analysis are aligned with previous reports of the ecological dynamics of the four insect groups selected in this study: cockroaches being present indoors all year round, with slightly higher frequency in warmer months, crickets chirping in late summer, mosquitoes with a steep peak over the summer^33^, and Monarch butterflies with a bimodal pattern with peaks in the spring and late summer, reflecting their well-known multigenerational migration from Mexico to Canada and back^28,34,35^.

The fact that similar patterns also emerge from the analysis of the specialized social network iNaturalist, where each entry most likely corresponds to a real observation, is also consistent with our hypothesis that the observed patterns in the Twitter data reflect phenological patterns.

Not only did our analysis allow us to identify periodic ecological patterns, but given the 11-year history of records, we could also observe changes in patterns over time. This, we argue, could be another advantage of using such sources of information: it may allow to study historical biological patterns retrospectively. These patterns could later be compared with climatic data to check for potential correlations and causal effects.

Our analyses, for example, show that tweets about mosquitoes have tended to extend longer in the season in more recent years, increasing in frequency in September and October. This pattern aligns with observations from New York state, where the season for West Nile Virus fever – a mosquito-borne disease transmitted by *Culex* species – has expanded over the past 25 years, starting 4 days earlier and extending 20 days further, likely due to climate change^36,37,38^.

However, for a more precise and reliable study of phenological pattern changes over time, future such studies should be conducted in a smaller area with a more homogeneous climate. In that case, changes in temporal patterns would be expected to correlate with local climatic data, helping to understand how changes in climate and environment are affecting different species.

### 3.2. Public perceptions, emotions, and conservation awareness

Our sentiment analysis based on the text of tweets suggest that social media may provide interesting insights into public perception of insect species, information that can be valuable for conservation campaigns, education activities, and policy making.

Disgust and fear were the predominantly negative emotions associated with cockroaches, consistent with existing stereotypes and psychological studies on animal fear and disgust^39^, while mosquitoes elicited anger, apparently because of the public health issues they pose^40^. Perhaps more surprising was the high proportion of “fear” related to the Monarch butterfly. The analysis of the most frequently used words in the text of the tweets where fear was the predominant feeling, revealed that this fear was related to the fear of extinction of this emblematic species. This highlights the increasing public awareness concerning the status of the species with regard to the decline of the overwintering populations and their populations generally^41,42^.

Our analysis of the most frequently used words in the tweet’s text also revealed additional information relevant to human-insect interactions, such as the most common locations and times where people found cockroaches (i.e. at night in the bathroom) or where they had most frequently been bitten by mosquitoes (i.e. forehead, face, and legs). We note that the information that can be retrieved from this analysis will be highly dependent on the query used. This caveat notwithstanding this analysis was useful to understand and explain the sentiment analysis results.

Overall, these analyses suggest that generalist social networks can offer a valuable window to understand societal perceptions of different species. Furthermore, we propose that this could also be used to track changes in perception over time. For example, it could be leveraged to evaluate the success of a given campaign about awareness of specific species in a specific region, comparing predominant sentiments before and after the given campaign.

### 3.3. Urban biases and spatial limitations

One of the aspects in which we hypothesized that social data could be very valuable, but failed to find supporting evidence for this hypothesis, was in providing spatial information that could help to track migrations or spread of invasive species.

A limiting factor of our Twitter dataset was the low proportion (∼3%) of geolocalized tweets, which might be expected to give the most direct evidence of the given observation. Even with this 3% of tweets, we were not able to obtain a clear pattern corresponding to known Monarch butterfly migration. We speculate that among the main issues for this was the lack of data for vast but poorly-populated areas^43,44^, and hotspots of activity around large metropolitan areas. On the other hand, the user location proved useful to filter tweets likely originating from a given country (e.g. USA and Canada). However this information was not precise enough to reveal the pattern of migration of Monarch butterflies.

While spatial monitoring was not supported by our dataset, we suggest that we should remain open to this possibility emerging from social media post in the future. For example there might be current or future networks that might geo-tag all posts, which could improve the resolution of this spatial monitoring.

### 3.4. Current and future directions for social networks in research

Since starting to collect the dataset for this study in 2022 and the time of writing (2025), many things have changed on Twitter. This platform, which once welcomed researchers to use their data, not only changed its name to X, but also imposed strong limitations on academic access in 2023^45^. As of the time of writing, there is no longer a free academic access tier such as the one we used herein. Based on the three current tiers ($0 USD/month 15,000 reads, $200 USD/month for 15,000 reads, and $5,000 USD/month for 1M reads) this project would require 2 months access to the highest tier and cost a minimum of $10,000 USD, which would make it economically unattainable for many laboratories.

Here, we used Twitter to provide evidence that a generalist social network could can provide valuable insights into ecology. But Twitter is not the only social network that exists, existed, or will exist. Thus, current (Facebook, Instagram, WeChat, Mastodon, Bluesky, Threads, TikTok, etc.) and future social networks may also have the potential to help scientists, not only in social sciences and epidemiology^46^, but as we show here, also in ecology and conservation biology. Given the high value of this information, we propose that social networks should establish mechanisms for scientists to access the data generated by the people to provide knowledge for the people.

We hope that with this work, reporting both the potential and the caveats of using social networks for research, we can encourage other researchers to use it for their studies. Here we also provide the technical framework, tools, and scripts (see this project’s GitHub) to facilitate these analyses.

We also recognize and admire the advantages of specialized social networks such as iNaturalist, which offer much higher-quality data for research. The main differences with the generalist social networks are the limited number of users and entries, the short temporal span as of the time of writing (although as shown in our analysis they have increased in the last 5 years), skewed entries towards certain regions and sociocultural backgrounds and reporting mostly species of scientific interest rather than casual observations.

While beyond the scope of this project, the analysis of images from photo-centric social networks like Instagram could provide an additional layer of interesting information through computer vision methods. Eventually, the integration of data from multiple platforms over the years might even help to provide a more comprehensive and complete picture of biological phenomena that society has passively accumulated.

## 4. Conclusions

- Using 11 years of Twitter posts for four groups of insects, we provide evidence that generalist social networks can provide temporal patterns that reflect insect biological patterns.
- This Twitter dataset also allowed us to study changes in the biological patterns over the years.
- Sentiment analysis of the tweets identified clear and distinct sentiments towards the queried insect groups.
- We have not been able to track geographical patterns over time, but based on our analysis, it seems plausible that this option could be possible if all posts were geotagged.
- We provide a methodological and conceptual framework for other scientists looking to use social data for biological research.

## 5. Methodology

### 5.1. Twitter data collection and preprocessing

We used the Twitter API v2^47^ for academic purposes (the project was submitted to and approved by Twitter, granting access to the Academic tier), to obtain tweets about monarch butterflies, crickets, mosquitoes, and cockroaches. The query included only tweets in English and excluded retweets and replies. The exact queries were as follows:

For for cockroach: “*(found OR saw) (cockroach OR cockroaches) -is:retweet -is:reply lang:en*”, for crickets “*(crickets OR cricket) chirp -is:retweet -is:reply lang:en*”, for monarch butterflies: “*monarch butterfly -is:retweet -is:reply lang:en*”, and for mosquito “*mosquito bite -is:retweet -is:reply lang:en*”.

In total, we downloaded 1.6 million tweets (downloaded: 29.01.2023), ranging between 62,289 and 1,043,571 per query (**Table 1),** which were used to plot the temporal pattern for each query in **Figure 1**. In the next step, we removed duplicated tweets, defined as those with identical text. Then, for tweets lacking geolocation data (namely, GPS coordinates), the user profile location field, if set up by the user, was extracted. The user profile location (usually the name of a place or city) was matched with OpenStreetMap^48^ API (v0.6) to approximate geographical coordinates, and these were assigned to the tweet. Temporal filtering was then applied to exclude tweets outside the 2011–2022 timeframe, ensuring the dataset was not influenced by the early years of Twitter, with a very limited number of posts. Additionally, content filtering was conducted to remove unrelated or spam entries, excluding tweets containing "etsy", "ebay", and "store". Following preprocessing, the final datasets comprised 36,413 tweets for cockroaches, 17,964 for crickets,145,528 Tweets for monarch butterflies, and 286,108 for mosquitoes (**Table 1**).

To identify seasonal trends across the dataset, we computed the number of tweets per day of the year for each year, while preserving their month and day attributes. Furthermore, we also computed the number of tweets from 11 years grouped by day of the year, day of the week, and time of day.

### 5.2. Temporal decomposition

The X11 model^49^, integrated within the X-13ARIMA-SEATS system developed by the U.S. Census Bureau, was employed using the seasonal^50^ package (v1.10.0) to decompose time-related data into key components, including trends and seasonal fluctuations.

The decomposition process began with seasonal adjustment, where the X11 algorithm iteratively estimated and removed recurring seasonal effects. This step isolated the trend-cycle and irregular components, enabling a clearer understanding of long-term patterns and irregular fluctuations. The trend-cycle analysis revealed gradual changes in tweet activity over time. Meanwhile, the seasonal component highlighted consistent intra-annual patterns.

To enhance interpretability, the decomposition results were visualized in R with ggplot2^51^ (v3.5.0) package. The same package was used for other graphical representations.

### 5.3. Seasonal changes over the years

To explore month-wise seasonal changes over the years, monthly tweet trends were extracted and analyzed for each taxa. By isolating data for each individual month, we captured detailed seasonal fluctuations unique to each month over time. These monthly time series were then aggregated and shown as seasonal amplitudes, enabling consistent cross-year comparisons.

### 5.4. iNaturalist data

Data from iNaturalist was downloaded from GBIF website (downloaded: 18.08.2023), selecting research-grade iNaturalist entries using as filter the following taxons Gryllidae for crickets, Blattodea for cockroaches, *Danaus plexippus* for Monarch butterfly, and Culicidae for mosquitoes. Additionally, we filtered entries from 2011-2022 for North America.

Time decomposition was performed for the Twitter dataset with the X11 method.

### 5.5. Sentiment analysis of tweets

We performed the sentiment analysis of the tweets using the Emotion English DistilRoBERTa-base^52^ model from Hugging Face ^31^, which is specifically trained for emotion recognition tasks in English. It of classifies text into emotion categories such as joy, fear, anger, disgust, surprise, sadness, and neutral.

With this model, we achieved a balance between computational efficiency and performance, making it suitable for large-scale analysis of social media data like tweets. This methodology was implemented in Python^53^ (v3.9.15) using the transformers^54^ (v 4.35.0) library from Hugging Face^31^. The model was applied without fine-tuning, using its general pre-trained capabilities to classify emotions effectively in social media text.

### 5.6. Spatial analysis

Geospatial analysis was performed to investigate the distribution of tweet activity across USA and Canada, providing insights into geographic trends and public interest hotspots. Tweets containing valid geolocation information (latitude and longitude) were selected from the dataset. These coordinates were rounded to the nearest degree to aggregate tweets into broader spatial bins, facilitating a clearer depiction of geographic trends. For visualization purposes, this was plotted as heatmaps of the density of tweets within these spatial bins, using R’s ggplot2^51^ package combined with the viridis^55^ (v0.6.5) library.

### 5.7. Ethical Considerations

The study complied with Twitter’s terms of service agreement for the academic tier and data usage policies, ensuring responsible handling and use of social media data. Only publicly available tweets were analyzed, and all data were anonymized to protect user privacy.

### 5.8. Code availability

The code used for obtaining the data and results is available at https://github.com/ylla-lab/Twitter-for-ecology.

## Author contributions

GY conceived of the project with support and feedback from CGE. RM performed the analysis and wrote an initial draft of the manuscript. GY wrote the final version of the draft, incorporating comments and suggestions from CGE. All authors read and approved the final version of this draft.

## Acknowledgments

This work has been supported by the Faculty of Biochemistry, Biophysics, and Biotechnology at Jagiellonian University (Poland) under the Strategic Programme Excellence Initiative (PRA BioS), the Federation of European Biochemical Societies Excellence Award 2023 (G.Y). C.G.E. is an investigator of the Howard Hughes Medical Institute.

