## Supplementary Figure for "Social media posts as a source of ecological information over time: using Twitter (X) as a proof of principle"

### Supplementary Figures

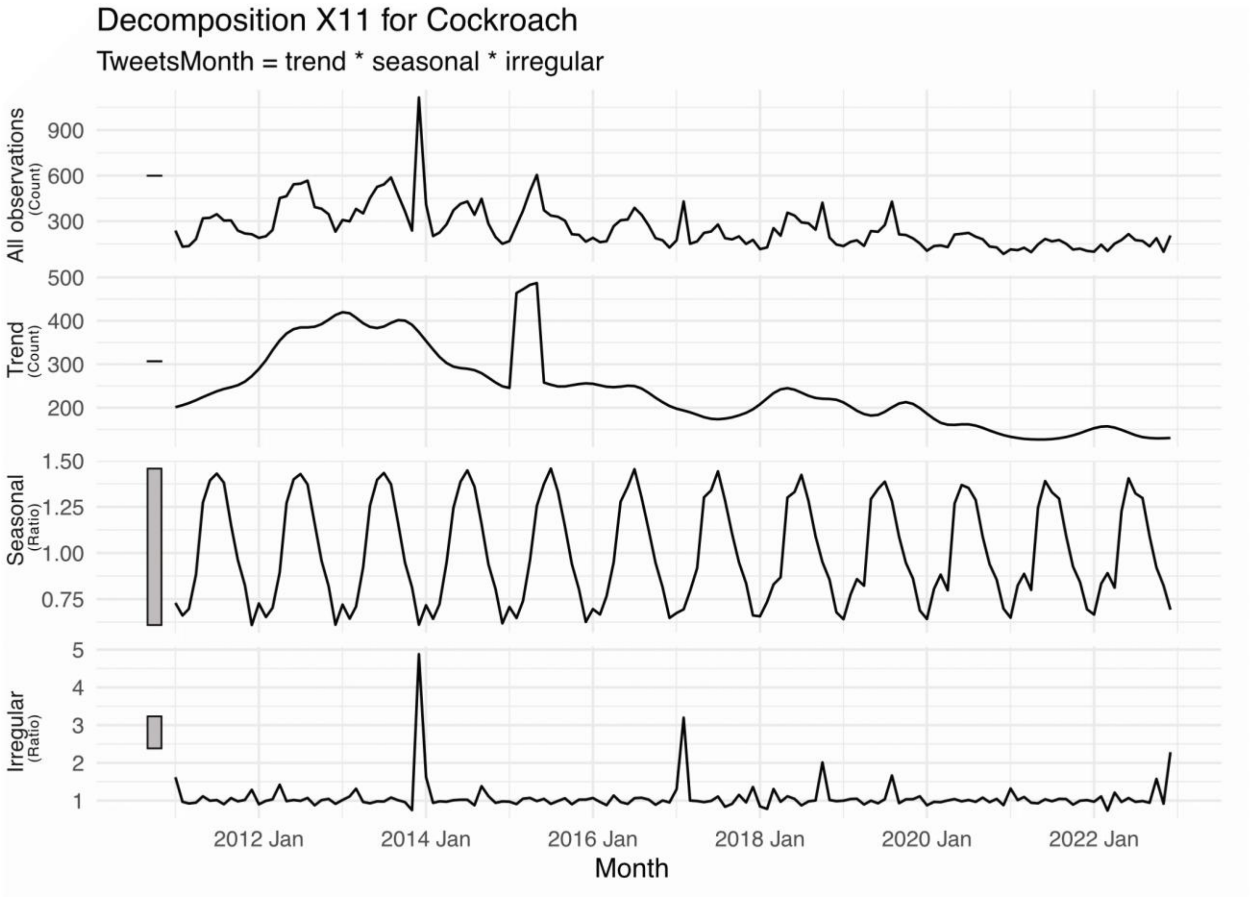


**Supplementary Figure S1:** Temporal decomposition with X11 method for the cockroach Twitter dataset into all tweets, trend, seasonal pattern, and irregular noise.

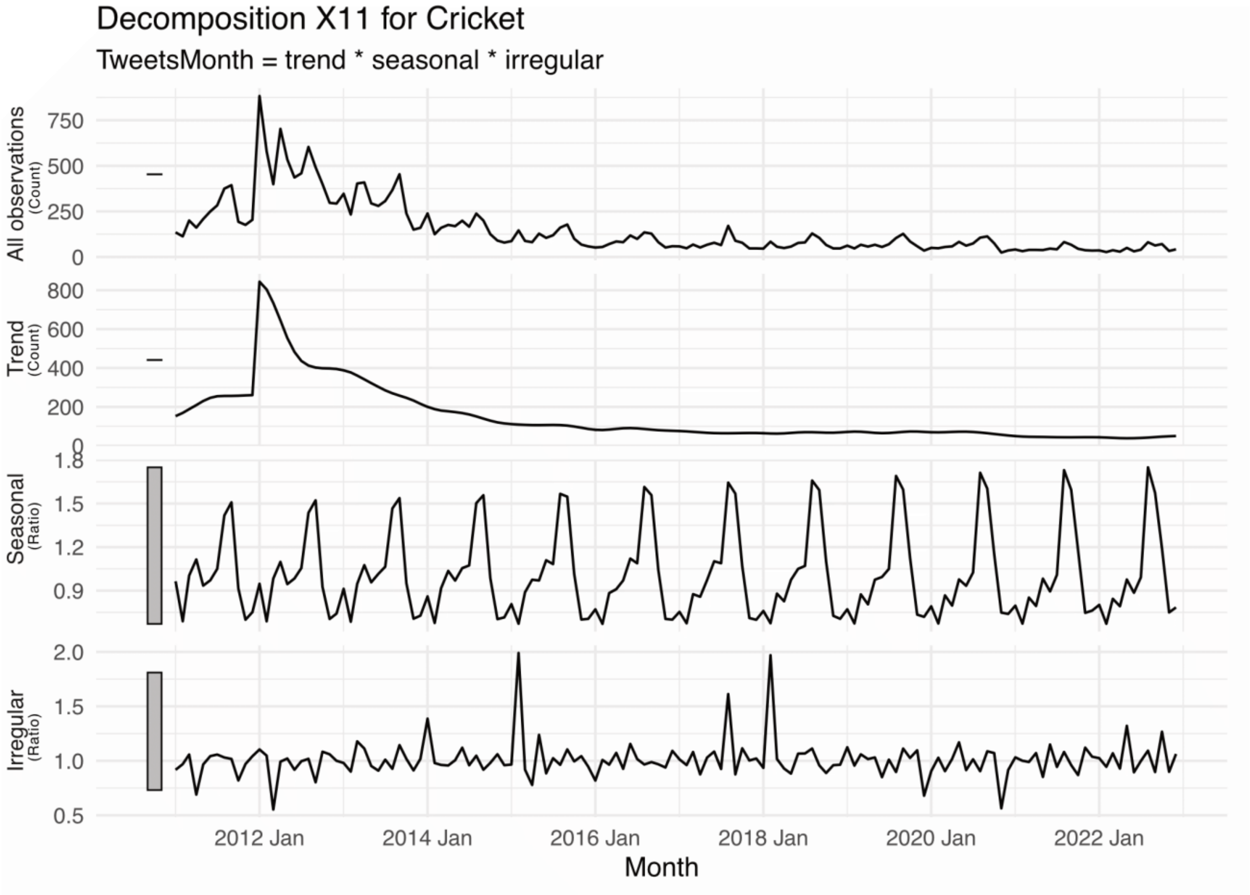


**Supplementary Figure S2:** Temporal decomposition with X11 method for the cricket Twitter dataset into all tweets, trend, seasonal pattern, and irregular noise.

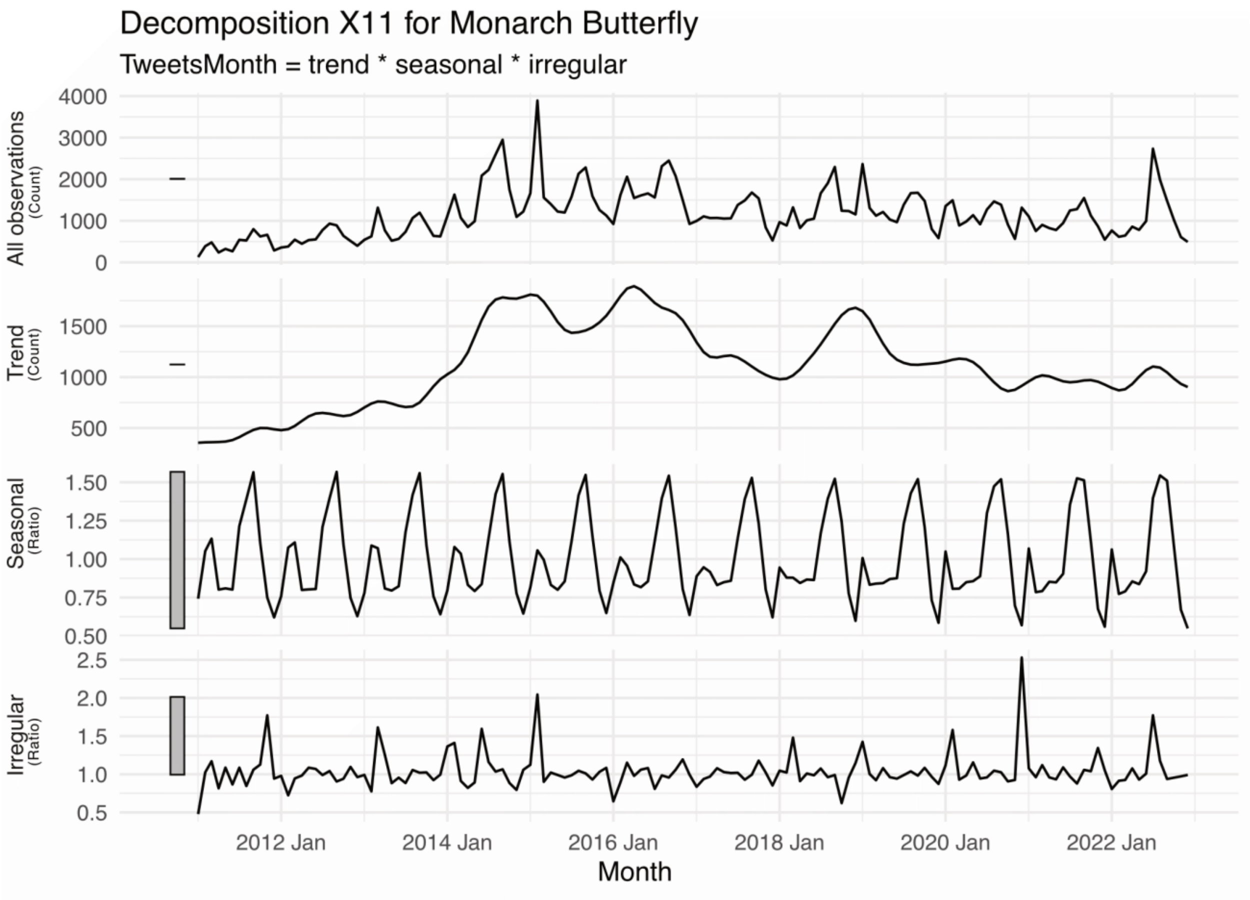


**Supplementary Figure S3:** Temporal decomposition with X11 method for the Monarch butterfly Twitter dataset into all tweets, trend, seasonal pattern, and irregular noise.

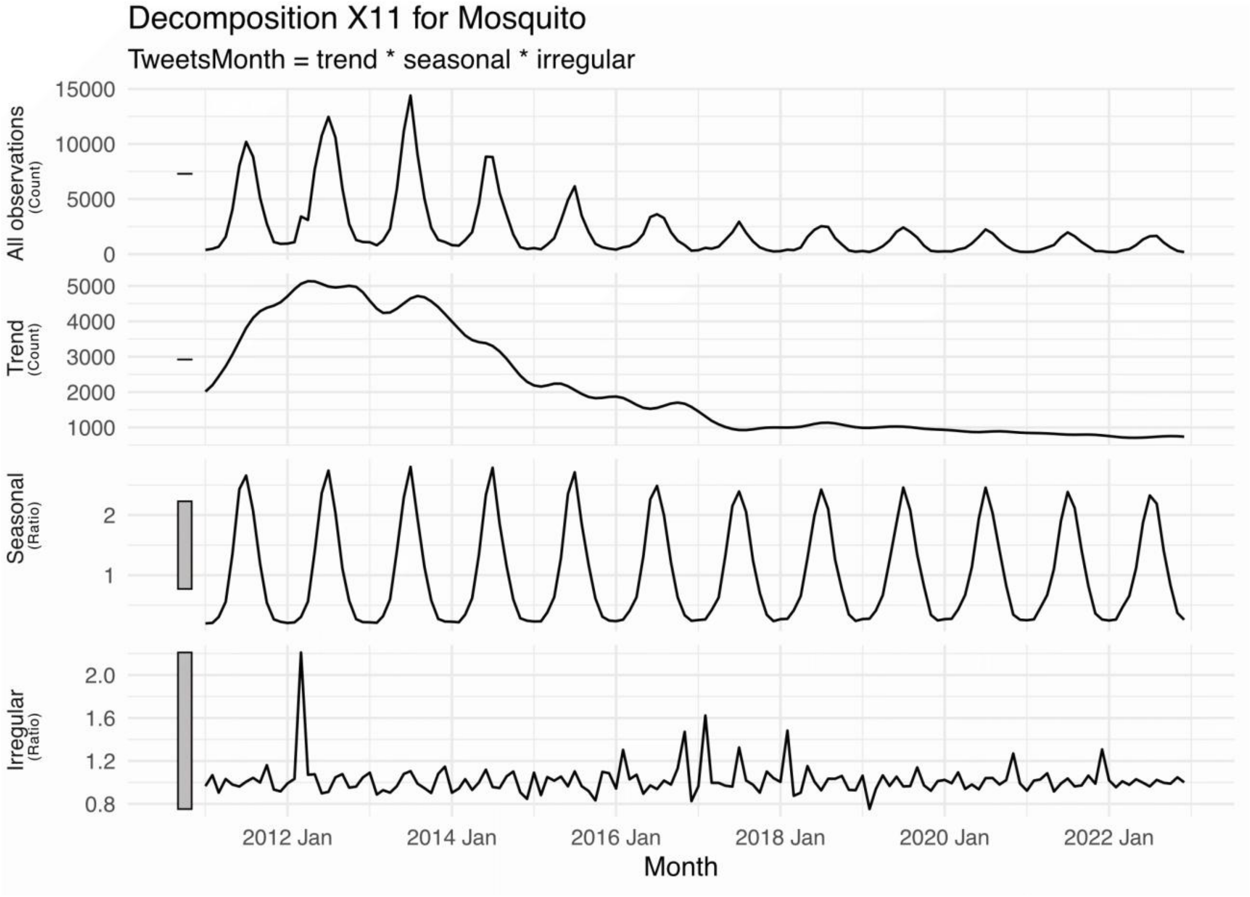


**Supplementary Figure S4:** Temporal decomposition with X11 method for the mosquito Twitter dataset into all tweets, trend, seasonal pattern, and irregular noise.

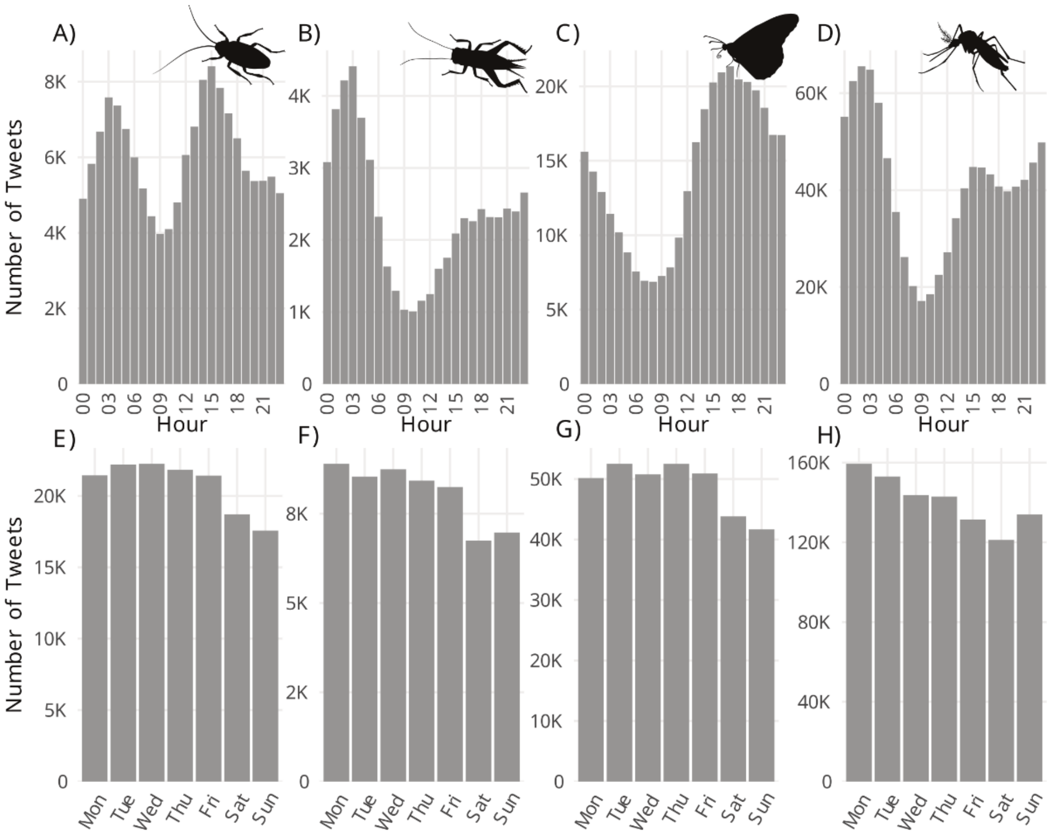


**Supplementary Figure S5**: Number of tweets for cockroaches (A & E), crickets (B & F), Monarch butterfly (C & G), and mosquitoes (D & H) per hour of the day (A to D) and by day of the week (E to H).

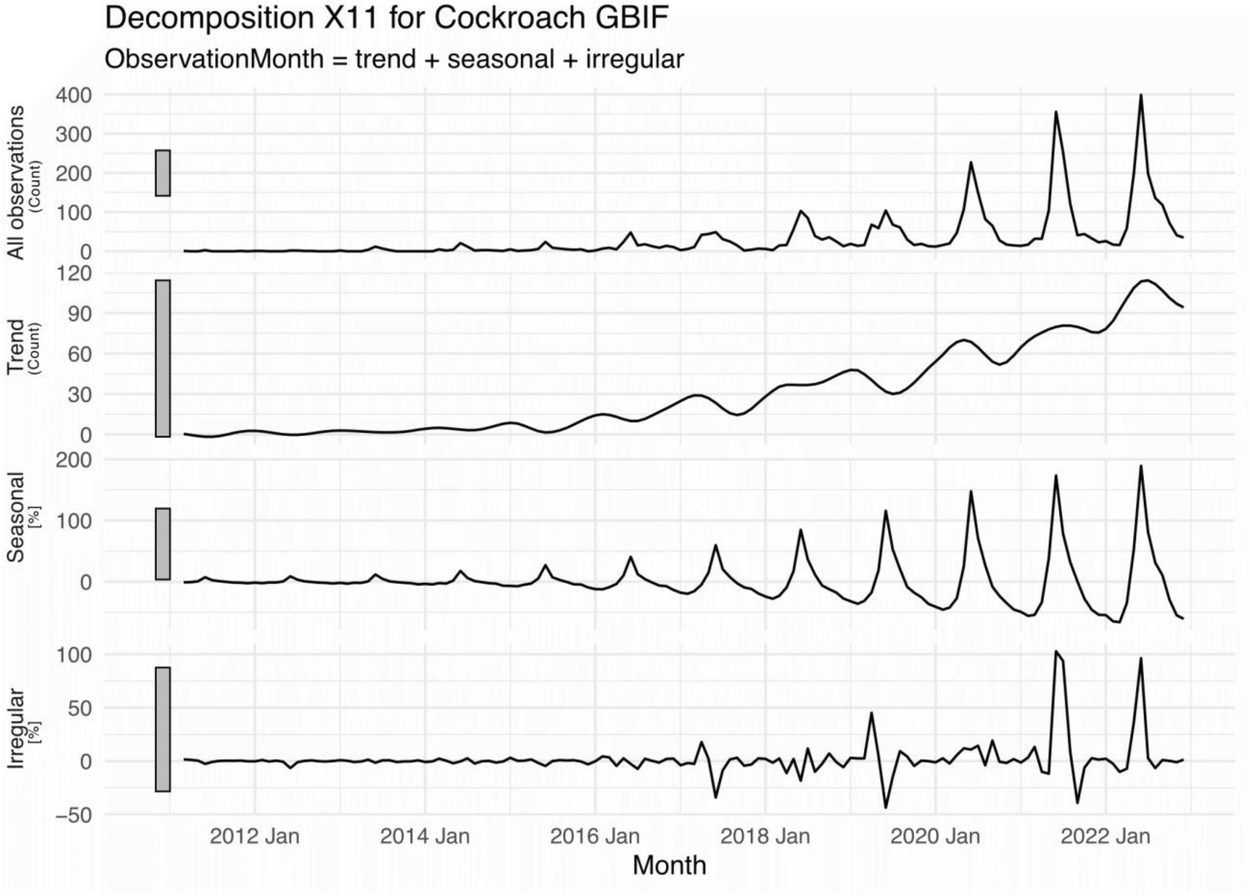


**Supplementary Figure S6:**Temporal decomposition with X11 method for the research grade entries for Blattodea in the iNaturalist dataset into all tweets, trend, seasonal pattern, and irregular noise.

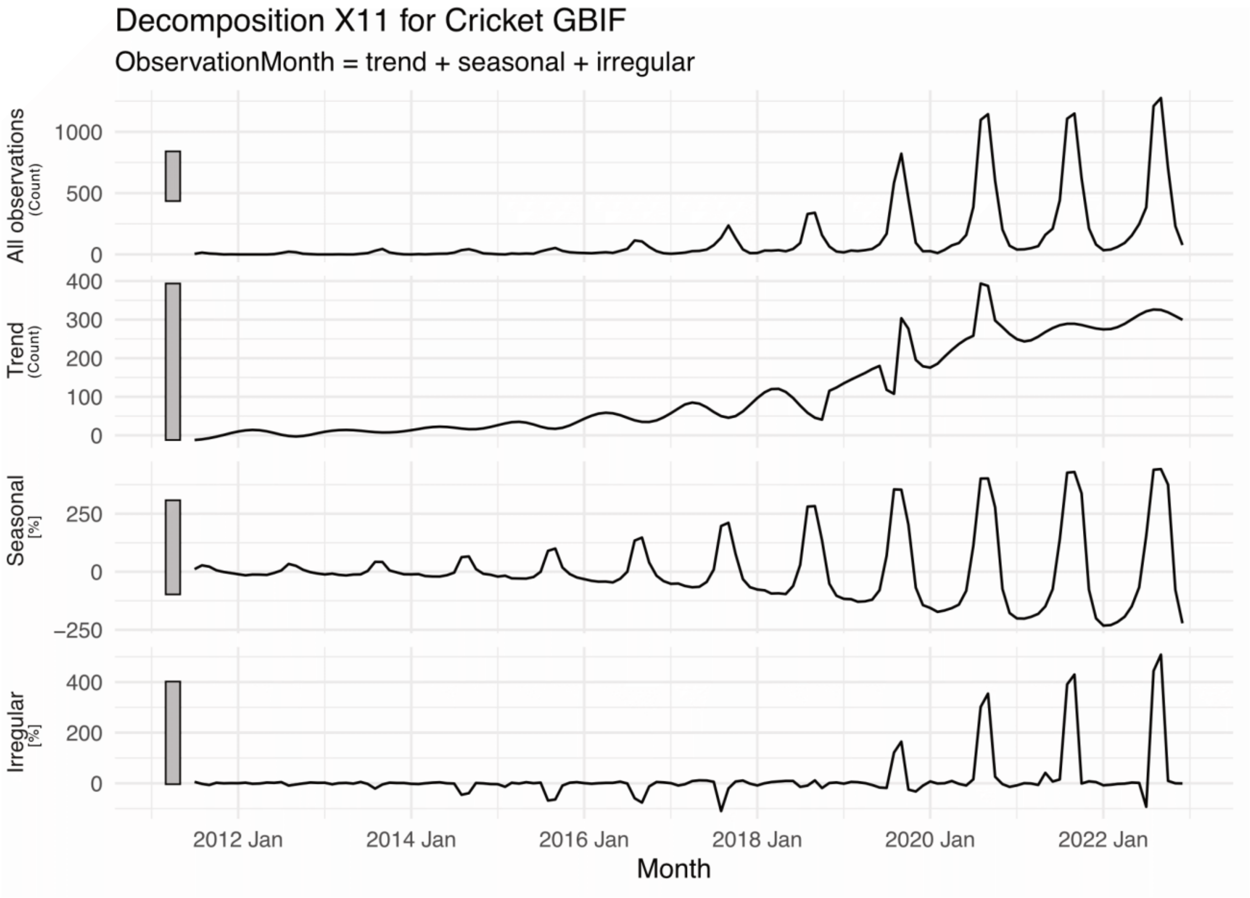


**Supplementary Figure S7:** Temporal decomposition with X11 method for the research grade entries for Gryllidae in the iNaturalist dataset into all tweets, trend, seasonal pattern, and irregular noise.

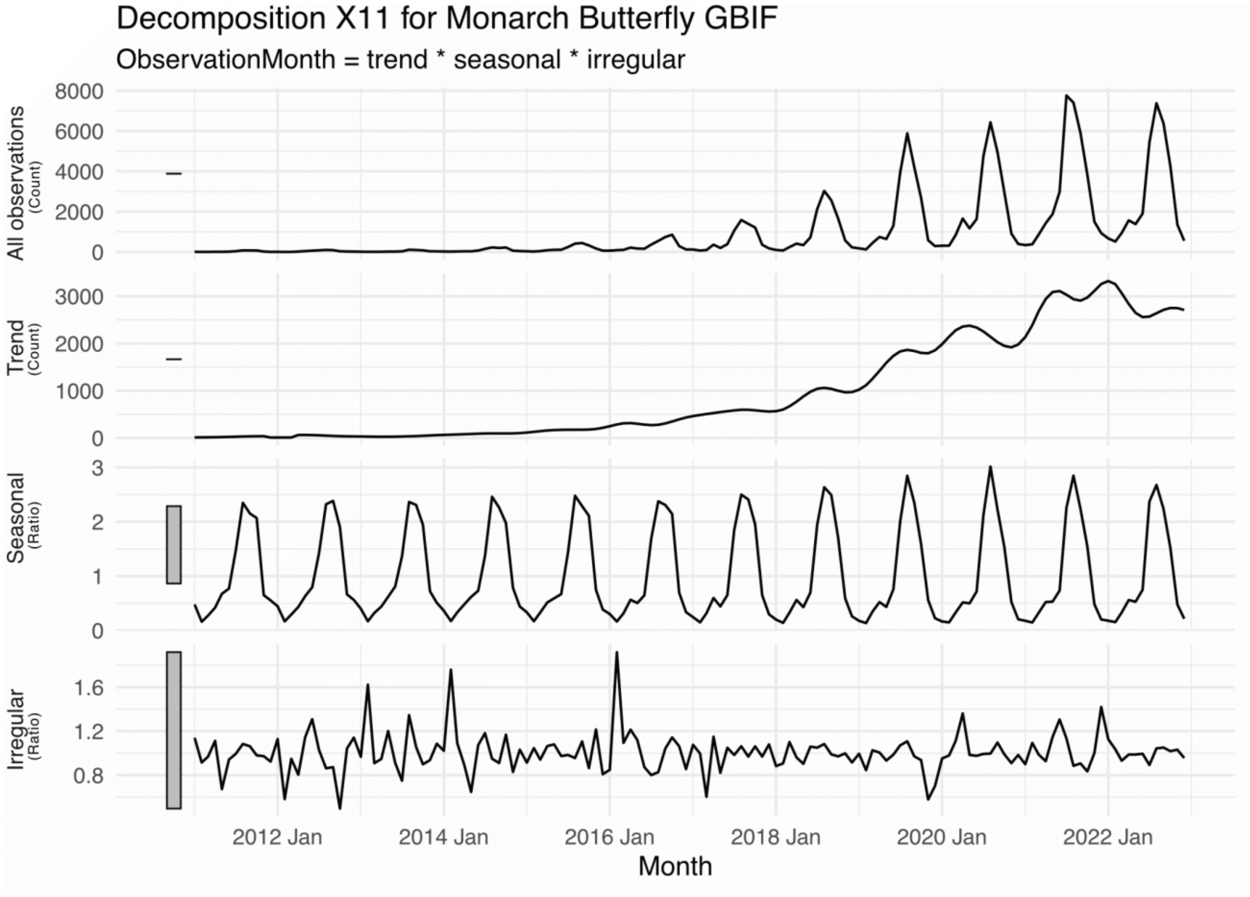


**Supplementary Figure S8:**Temporal decomposition with X11 method for the research grade entries for *Danaus plexippus* in the iNaturalist dataset into all tweets, trend, seasonal pattern, and irregular noise.

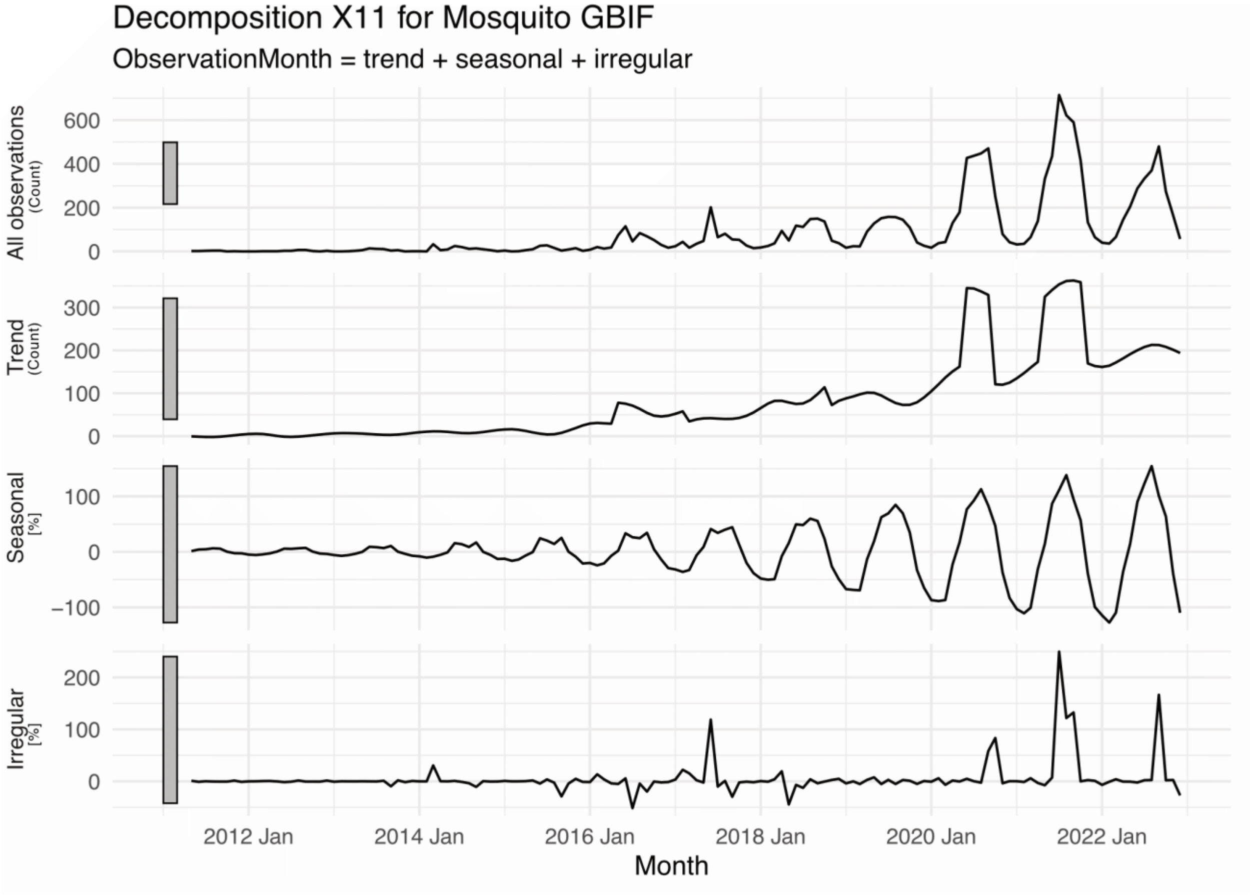


**Supplementary Figure S9:**Temporal decomposition with X11 method for the research grade entries for Culicidae in the iNaturalist dataset into all tweets, trend, seasonal pattern, and irregular noise.

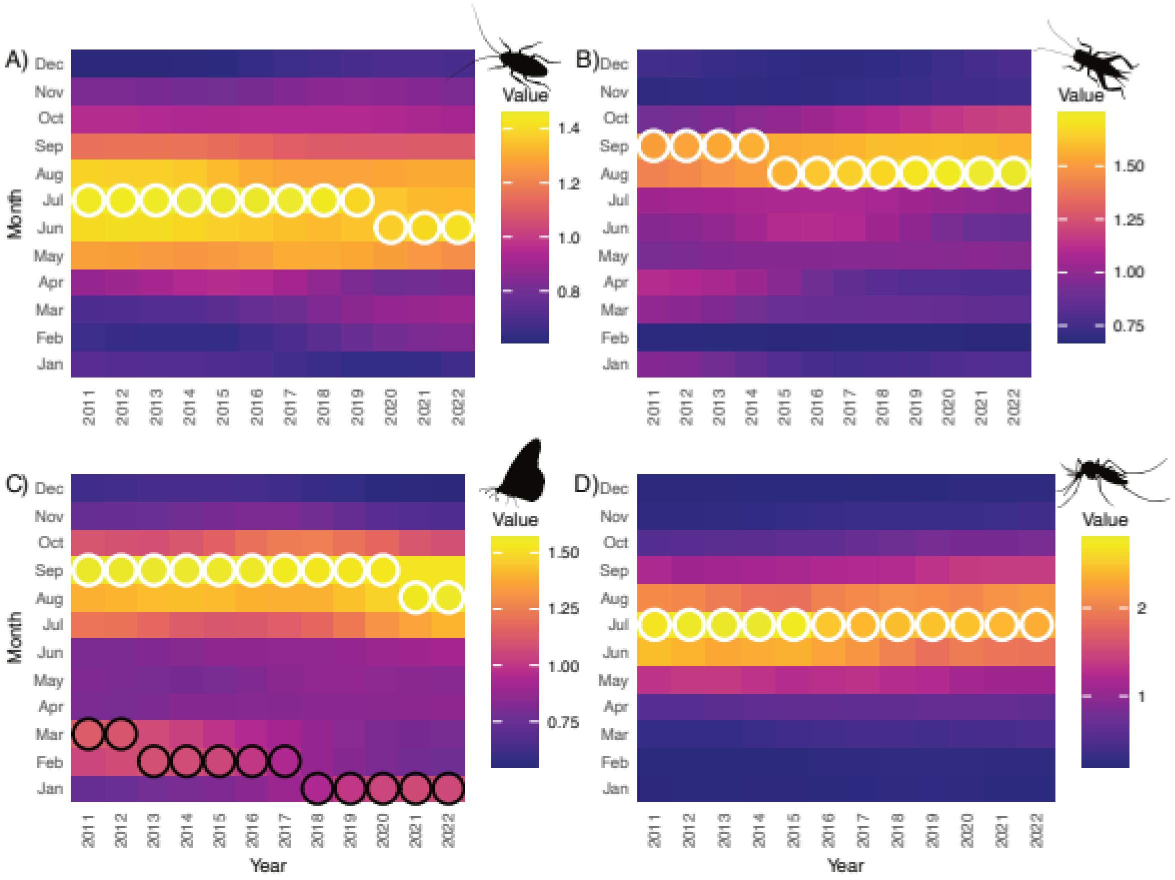


**Supplementary Figure S10**: Heatmap of monthly activity by calendar year for each dataset. For each taxa and year, the primary annual peak (absolute maximum) is indicated by a white open circle. For Monarch Butterfly only, a black circle marks the second within-year peak defined as the last local maximum occurring prior to the annual maximum.

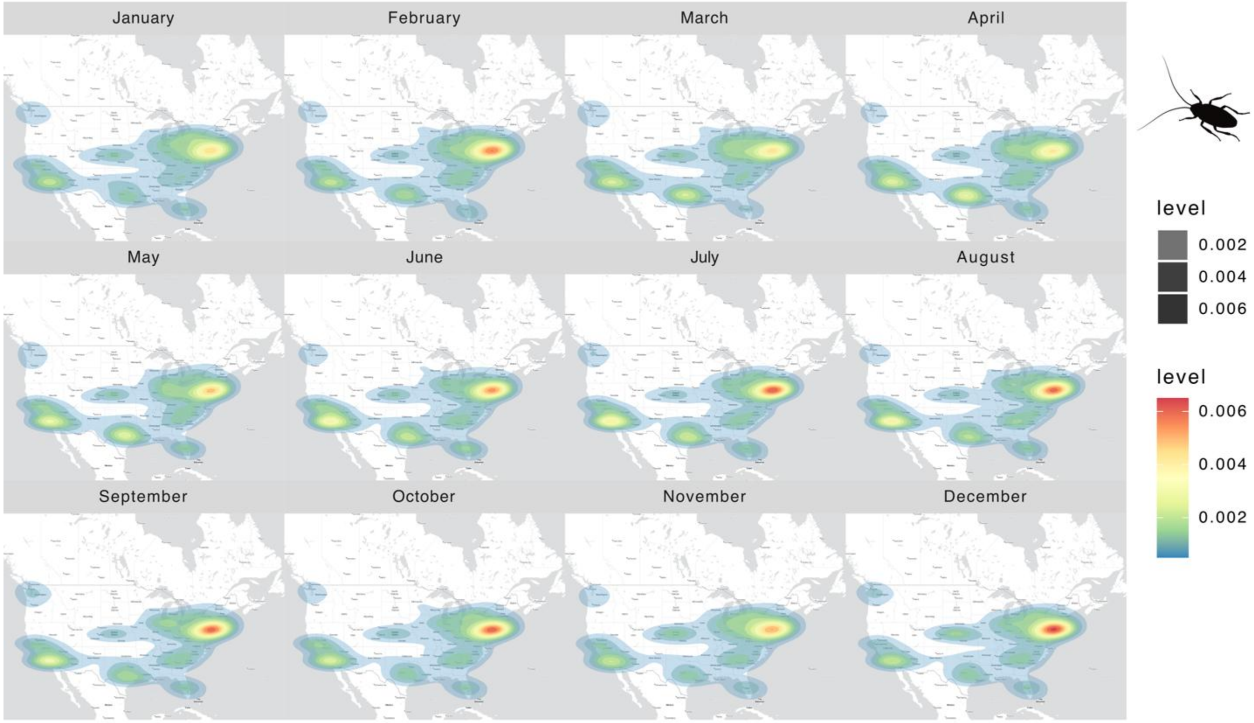


**Supplementary Figure S11: Monthly spatial intensity of geolocated cockroach-related tweets.** Each panel shows a calendar month (Jan–Dec); filled polygons depict a 2-D kernel density estimate of tweet locations (higher “level” = higher relative intensity). Color and opacity scale with the estimated density; panels share the same geographic extent to allow visual comparison across months.

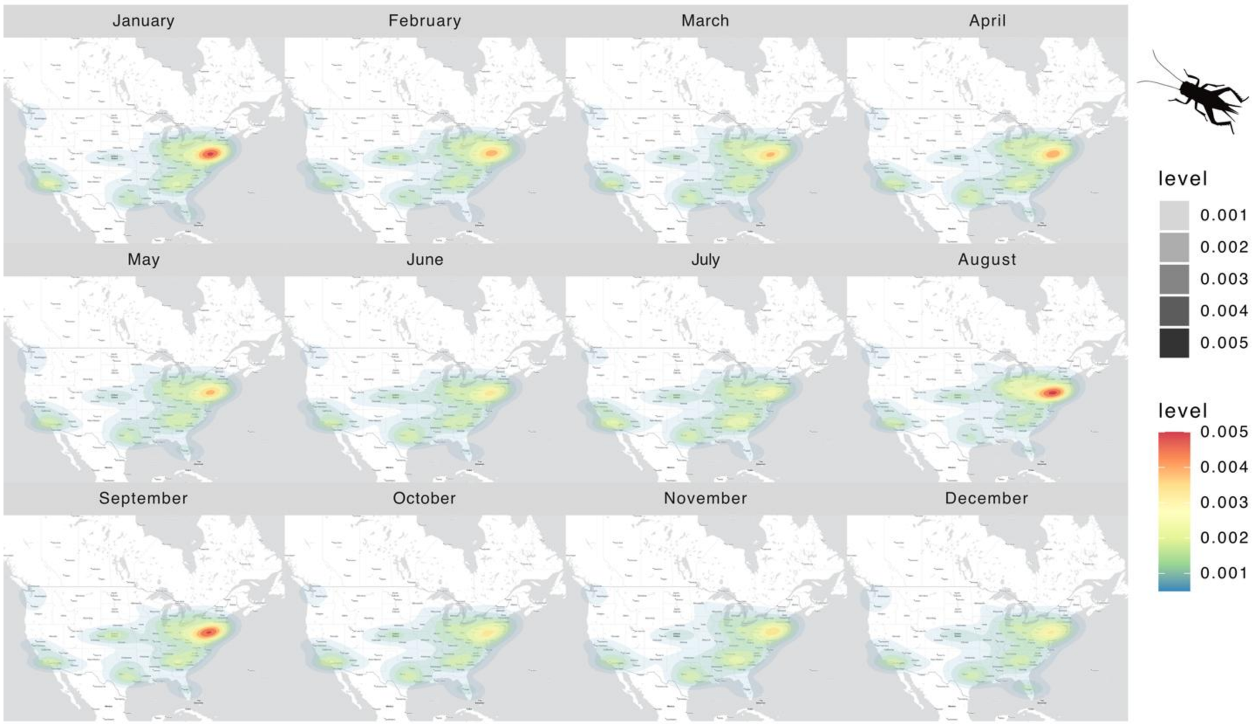


**Supplementary Figure S12: Monthly spatial intensity of geolocated cricket-related tweets.** Each panel shows a calendar month (Jan–Dec); filled polygons depict a 2-D kernel density estimate of tweet locations (higher “level” = higher relative intensity). Color and opacity scale with the estimated density; panels share the same geographic extent to allow visual comparison across months.

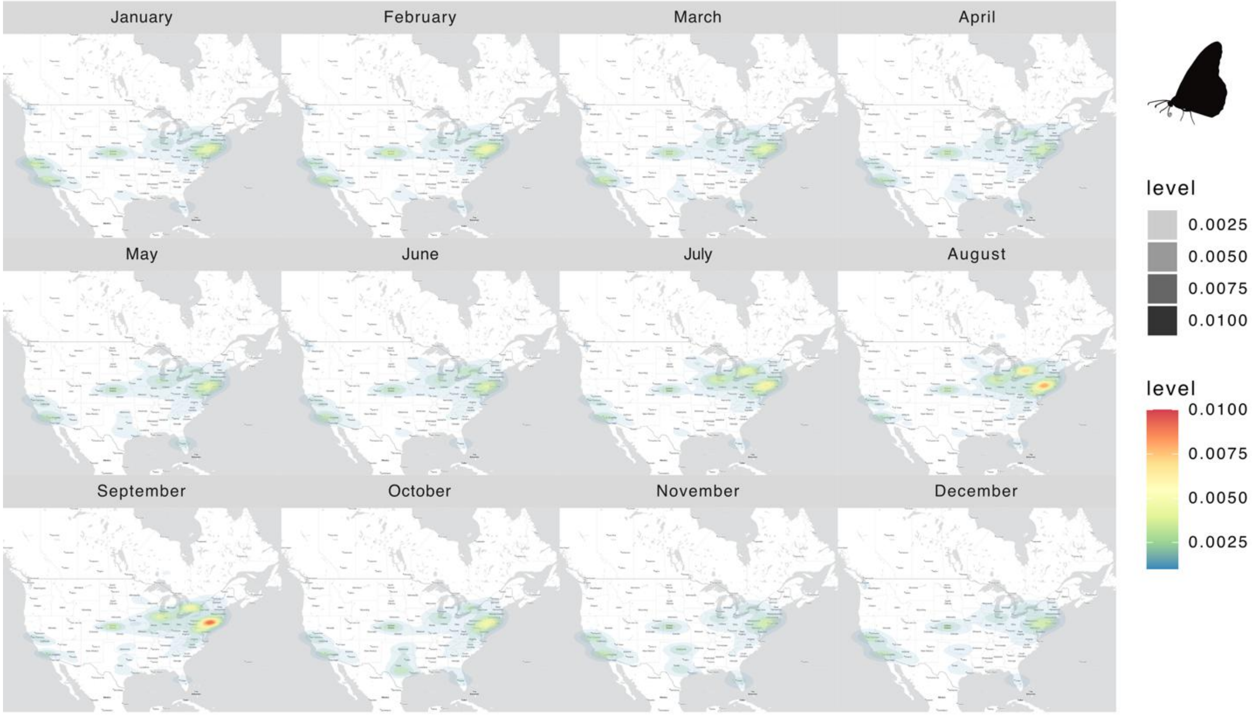


**Supplementary Figure S13: Monthly spatial intensity of geolocated Monarch-related tweets.** Each panel shows a calendar month (Jan–Dec); filled polygons depict a 2-D kernel density estimate of tweet locations (higher "level" = higher relative intensity). Color and opacity scale with the estimated density; panels share the same geographic extent to allow visual comparison across months.

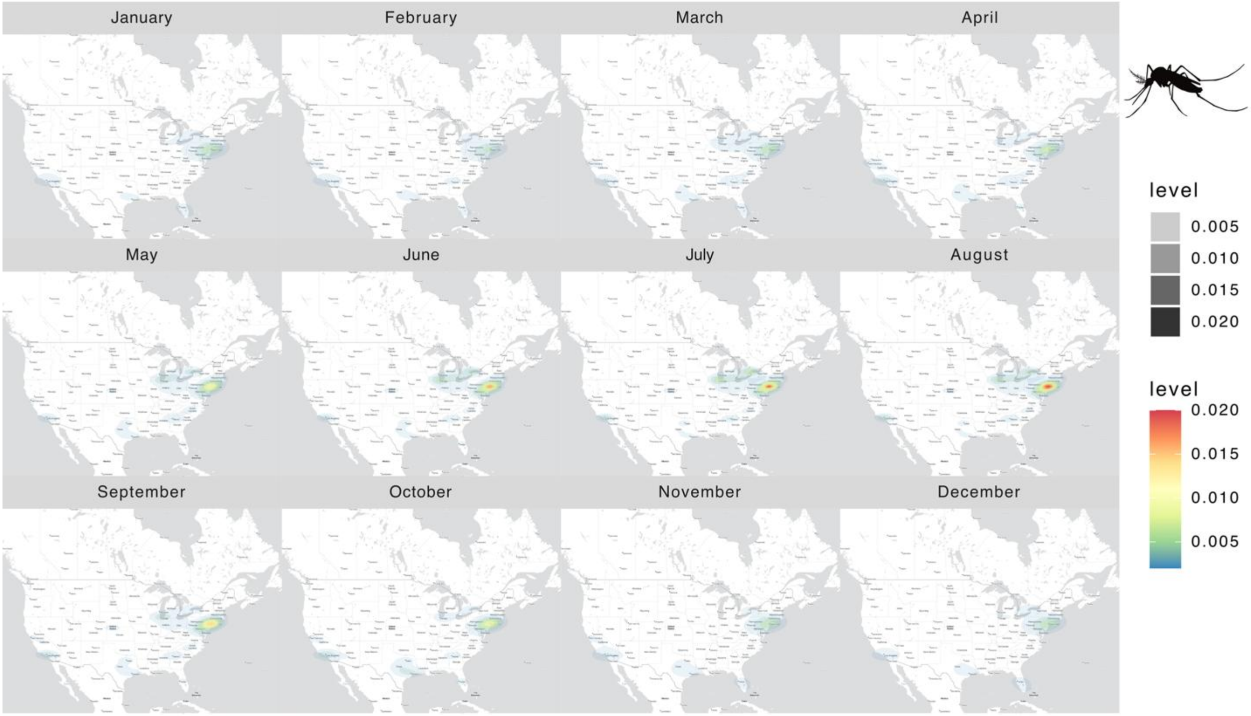


**Supplementary Figure S14: Monthly spatial intensity of geolocated mosquito-related tweets.** Each panel shows a calendar month (Jan–Dec); filled polygons depict a 2-D kernel density estimate of tweet locations (higher “level” = higher relative intensity). Color and opacity scale with the estimated density; panels share the same geographic extent to allow visual comparison across months.
